# Interpersonal physiological synchrony: estimation and clinical application to cardiac dynamics of parent-infant dyads^⋆^

**DOI:** 10.64898/2026.03.19.712915

**Authors:** Laura Lavezzo, Didier Grandjean, Sylvain Delplanque, Francisca Barcos-Munoz, Cristina Borradori-Tolsa, Enzo Pasquale Scilingo, Manuela Filippa, Mimma Nardelli

**Affiliations:** Bioengineering and Robotics Research Centre E. Piaggio and Dipartimento di Ingegneria dell’Informazione, University of Pisa, Largo Lucio Lazzarino 1, Pisa, 56122, Italy; Department of Psychology and Educational Sciences and Swiss Center for Affective Sciences (CISA), Campus Biotech, University of Geneva, Chemin des Mines 9, Geneva, 1202, Switzerland; Division of Pediatric Intensive Care and Neonatology, Department of Women, Children and Adolescents, University Hospital of Geneva, Geneva, 1205, Switzerland; Division of Development and Growth, Department of Pediatrics, University Hospital of Geneva, Geneva, 1205, Switzerland

**Keywords:** Physiological synchrony, Heart rate variability, Synthetic series, Premature infant, Autonomic nervous system, Multivariate multiscale entropy, Complexity, Coupling

## Abstract

Synchrony is a key mechanism that builds up the foundations of human interactions. Quantifying the level of physiological synchronization that occurs during dyadic exchanges is essential to fully comprehend social phenomena. We present a new index to characterize the coupling of complex physiological dynamics: the optimized Multichannel Complexity Index (*op*MCI). We validated this approach using synthetic time series of two coupled Hénon Maps, with four different coupling levels in unidirectional and bidirectional manners. We demonstrated that the *op*MCI method allows to effectively discern between all coupling levels. Then, we applied the *op*MCI metric on heart rate variability data collected from 37 parent-infant dyads, during shared reading and playing activities, in the framework of the Shared Emotional Reading (SHER) project, with the aim of assessing the effects of early intervention in preterm babies. Two groups presented preterm infants: an intervention group, who participated in a two-month shared reading program, and a control group, who practiced shared play activities. A full-term group provided additional control data. The *op*MCI values were significantly higher for the intervention dyads with respect to the other groups during the shared reading task, showing that an early reading intervention program could increase parent-infant synchrony in preterm babies.

**Graphical Abstract:** 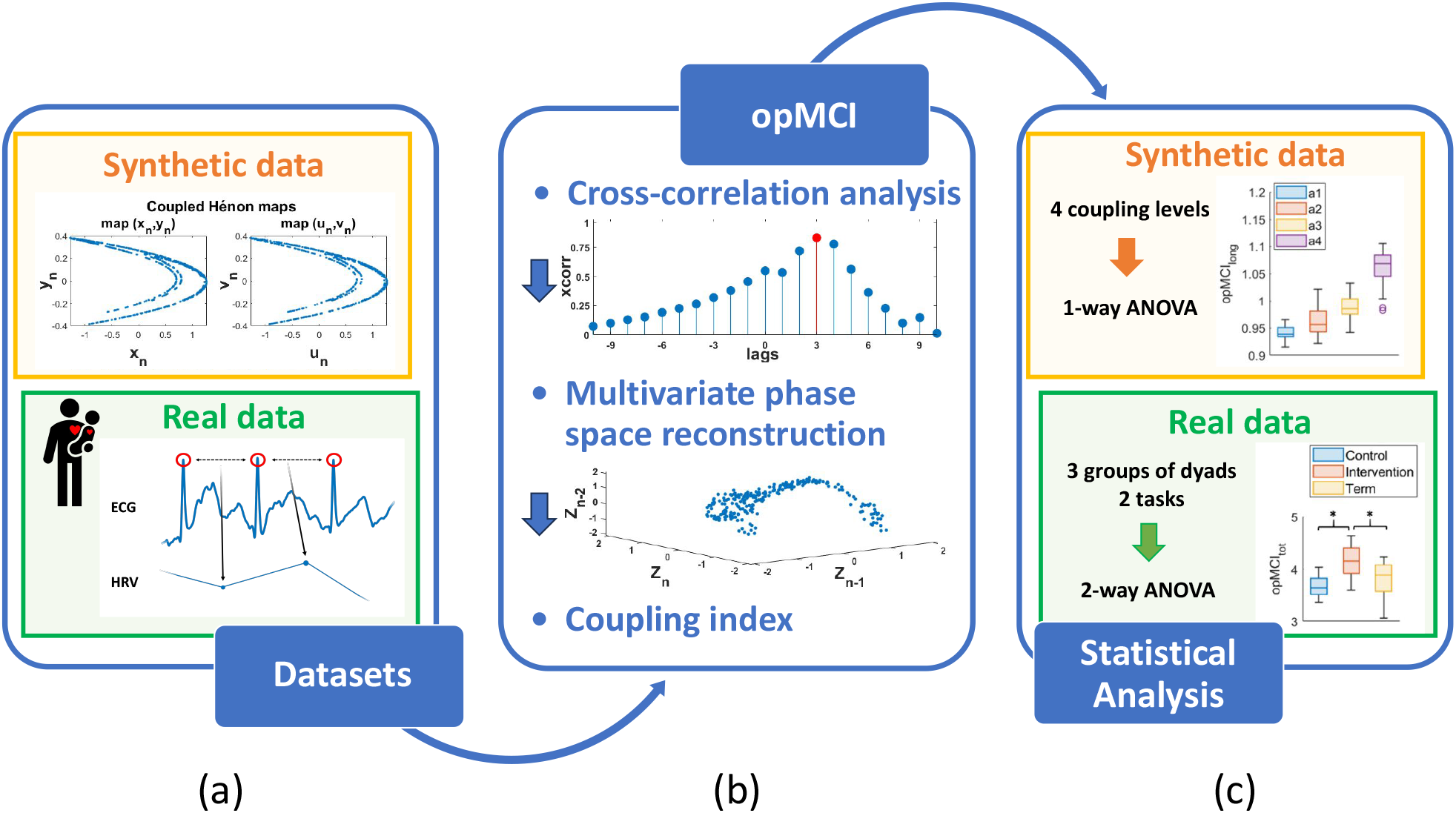

**Highlights:**

- Interpersonal physiological synchrony is a fundamental mechanism for the evolutionary path
- An ad-hoc methodological framework to assess nonlinear cardiac coupling is lacking
- The *op*MCI has been shown to be a reliable tool for assessing the degree of synchrony
- Early shared book reading may increase parent-infant synchrony in a clinical setting

## 1. Introduction

Human physiological systems are extremely responsive to their sensory environment, able not only to dynamically detect changes, but also to adjust their processes accordingly, even while stimulation is occurring (1). In this context, a particularly refined human ability is to frequently and effortlessly synchronize behavioral and physiological dynamics with other individuals (2), a fundamental mechanism for the evolutionary path that supports the development of social communities (3). Social synchrony leads to several benefits, such as the facilitation of co-regulation and mutual adjustment of physiological processes (4), and the enhancement of cooperation (5) and cohesion within groups (6; 7; 8). Specifically, previous literature showed that physiological synchrony has been found to be a predictor of both trust (9) and compassion (10), and is a cornerstone for the development of bonds and social attachments, playing a key role in the formation of human relationships (11; 12).

The parent-child dyad is considered the gold standard social relationship for synchrony-related studies (13). Several previous studies have shown a significant positive association between the parent-child synchrony and children’s developmental outcome, in particular in the social, emotional and cognitive domains (14; 1; 15). Parent-infant synchronization is at the base of infant-caregiver coregulation of affective states (16; 17; 18; 15), it facilitates a secure attachment (17), and it is functional to language acquisition (18). Furthermore, synchrony plays a crucial role in the development of infant self-regulation, a protective developmental function that allows infants to dynamically adjust their physiological responses to the environmental requests(1; 16; 19). Parent-infant synchronization is especially compromised in high-risk populations, such as premature babies. Preterm infants show an atypical maturation of the autonomic nervous system (ANS), with short- and long-term related adverse impacts (20; 21; 22). Preterm birth is also known to have negative impact on parental sensitivity levels towards their infants and on parental stress and depression levels (23; 24). As a result of a multiple array of causes, premature infants show lower synchronicity during interactions with their parents, while they would benefit the most from it (14; 24).

A physiological quantification of parent-infant synchrony can be useful for both gaining a more specific understanding of its influence on the developmental process and assessing the potential beneficial effect of early interventions on the dyad. Considering the specific high-risk population of preterm infants, the study of ANS dynamics represents an optimal target for the assessment of dyadic synchrony and infant neural maturation, also allowing a non-invasive monitoring compatible with the performance of relational-based tasks.

Human body rhythms arise from nonlinear systems and are the result of stochastic interaction between complex physiological mechanisms and a fluctuating environment (25). The commonly used approach in literature is the conceptualization of the cardiovascular system of an individual as a network of biological self-sustained oscillators, with feedback loops reacting to internal and external inputs. The system’s dynamics can be revealed through the analysis of the time course of a systemic variable (26). Consequently, techniques derived from chaos theory, which consider inherent nonlinearities, are necessary for accurately and comprehensively describing the physiological dynamics, capturing the irregular fluctuations and helping to manage potential pathological rhythms (25). During the last few decades, these methodologies have shown promising results, in particular in the field of emotion recognition (27; 28) and mood disorder assessment (29; 30). Extending these results from the individual characterization to dyads and group dynamics could provide key insights into the synchronization phenomena. A conceptualization based on complex dynamical systems was previously introduced to model human social collective behaviour as well, with humans as sub-systems of a social unit capable of exchanging information through interactions (31; 32), with physiological aspects related to synchronization recently added into the picture (33). Based on this description of the social interaction phenomena, considering the complex dynamics present at both individual and interpersonal levels, we chose to focus on chaos-based techniques to model physiological synchrony. To this aim we propose a novel approach, i.e. the optimized-Multichannel Complexity Index (*op*MCI), based on chaos theory, to analyze physiological synchrony. The algorithm consists in an optimization of a previously developed method (34) for the computation of the complexity of nonlinear multivariate processes, here extended to account for the timing of social dynamic interactions. The optimized and the standard MCI approaches have been tested on the time series obtained from both coupled synthetic systems (coupled Hénon maps) and real-cardiovascular data recorded from parent-infant dyads. Specifically, we assessed the cardiovascular synchrony level in dyads composed of term or preterm infants with their respective parents, during shared tasks based on play and reading activities.

## Methods

All analyses have been performed using Matlab 2021b unless otherwise specified.

### Synchronization metrics

#### The Multichannel Complexity Index (MCI)

The Multichannel Complexity Index (MCI) (34) is a tool to characterize the complexity and coupling of multivariate processes.

According to the method described in (34), the MCI is obtained through a multiscale approach, that reconstructs *β* coarse-grained series from the original time-series by computing the average within non-overlapping windows of *β* data points. Considering 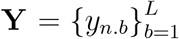 as a c-variate time-series with *n* = 1, …, *c*, of length *L*, each element of the coarse-grained series 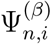 at scale *β* can be computed as:

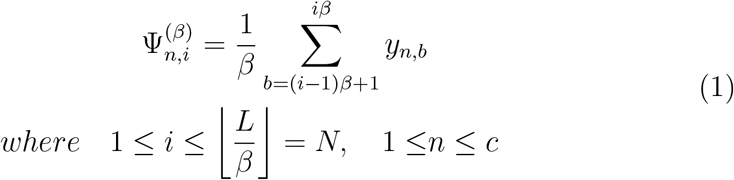

In previous literature (35; 36), the range of the scale factor *β* was reported to be related to the observation of different phenomena:

1. *β <* 5 indicates the short range scales, depicting fast oscillations phenomena
2. *β >* 5 reports on the long range scales, depicting slow oscillations phenomena

Limited data-length can affect the reliability at higher scales, to account for this the long range in this work was limited to *β* ∈ [6, 10].

For each coarse-grained series a c-variate phase space is computed, defining first the embedding dimensions for each channel [*m*_1_, *m*_2_, …, *m_c_*], and the time-delays [*τ*_1_, *τ*_2_, …, *τ_c_*]).

The reconstruction of the multivariate phase space is performed through a median procedure between all the trajectories of the single channels, obtaining the multivariate embedded vectors *Z*_**M**_ as follows:

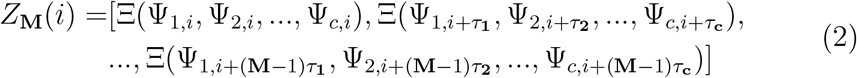

with Ξ representing the median value of the samples, and **M** = max ([*m*_1_, *m*_2_, …, *m_c_*]) is the dimension of the c-variate phase space.

Then the values of the Chebyshev distances *d* between pairs of embedded vectors are weighted using the fuzzy membership function 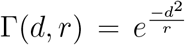, where 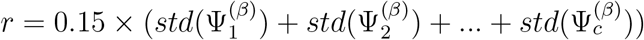 (37; 38).

The average membership degree is computed as:

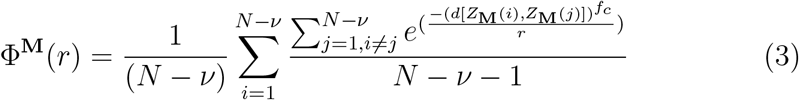

where **T** = *max*([*τ*_1_, *τ*_2_, …, *τ_c_*]) and *ν* = **T** × **M**. Finally, the multivariate entropy metrics for each value of *β* is:

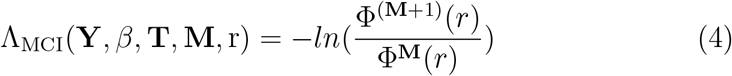

Following this procedure, a multiscale trend of the entropy level as a function of the scale factors *β* is obtained. From these trends two MCI values are defined as the area under the curve for the short- *β* ∈ [1, 5] and the long-range *β* ∈ [6, 10] scales (MCI*_short_* and MCI*_long_*, respectively).

In (34) a statistical significant difference was found between the MCI values of synthetic multi-channel systems with different inter-channel coupling levels. Expanding on these findings, the capability of the MCI metrics to discern between different levels of coupling has been further tested on both coupled synthetic systems and physiological data of parent-child dyads.

#### The proposed optimized-MCI (opMCI)

A novel approach, i.e. *op*MCI, is here proposed to optimize the MCI technique with the aim to improve performance in identifying the nuances of nonlinear synchronization between time-varying processes. A new step is introduced at the beginning of the MCI algorithm, and is based on the calculation of the maximum positive peak of the cross-correlation function between the time series involved. Given a bivariate process of L samples, 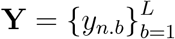 with *n* = 1, 2, we calculate the following value:

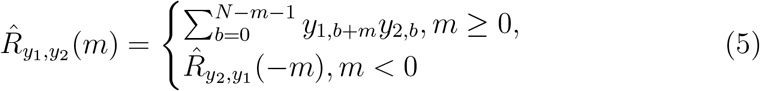

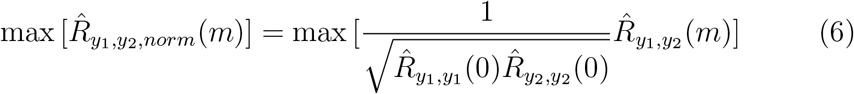

Therefore, we apply the MCI algorithm in a new optimized configuration of synchrony, and we compute the value of *op*MCI by shifting the two input time series for an amount of samples corresponding to the lag *m* of the maximum cross-correlation coefficient. The realignment procedure could provide a more comprehensive assessment of the phenomenon, considering both the strength of the coupling and possible time-delayed effects. These two factors are examined sequentially, first considering the lag through the cross-correlation approach, then assessing the coupling strength after removing the influence of the lag through the realignment. Analyzing first the lag itself could provide a quick and easy-to understand metric to quantify the delays in the couple rhythms. While the analysis of the realigned signals would allow to examine the strength of the coupling in an “optimal framework”.

#### Experimental Data

The MCI and *op*MCI metrics were extracted from both coupled synthetic dynamical systems and physiological data of parent-child dyads during interacting tasks. Details on the two datasets follow below.

#### Synthetic data

The *op*MCI has been tested on two coupled Hénon maps, to consider a synthetic dynamical process as a model of synchrony phenomena(39; 40):

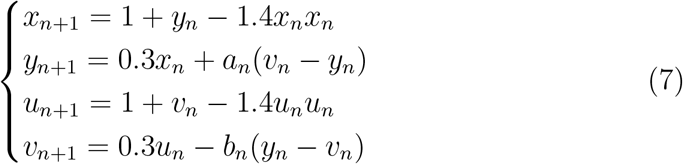

The coupling between the two subsystem (*x_n_, y_n_*) and (*u_n_, v_n_*) is determined by the values of the parameters *a_n_* and *b_n_*. Since synchrony in a multivariate process is a transient phenomenon (17) where the subsystems move closer or further away from a state of coupling during the interaction, the parameters can vary to mimic a change in synchronization dynamics. The synchrony level between the time-series *x* and *u* has been assessed in different configurations with increasing level of coupling. For each pair of coupling parameter (*a_n_,b_n_*), we generated 50 realizations of 5000 points each, with additive white gaussian noise of amplitude 0.1 added afterwards. We studied two different configurations of synchrony: unidirectional and bidirectional coupling. The first configuration was obtained by fixing *b_n_* = 0 during the whole simulation, and changing *a_n_* from 0 to a fixed threshold, after a number of samples randomly selected in the interval [100 − 300], to mimic the physiological synchronization behaviour. We investigated four values of *a_n_*, i.e., [0, 0.02, 0.04, 0.08], according to (39). The second configuration was tested with the same condition for *a_n_*, and setting the value of *b_n_* to 0.03 (40).

The MCI and *op*MCI metrics were used to discriminate between different levels of coupling parameter *a_n_* in each of the two configurations.

#### Physiological data

We evaluated the performance of MCI and *op*MCI for the assessment of cardiovascular synchrony during the social interaction of two individuals. Specifically, we investigated the parent-infant dyad, as the gold-standard for dyadic interaction and physio-behavioural synchronization studies.

We analysed the cardiovascular data of 37 dyads acquired during a preliminary study in the framework of the research project Shared Emotional Reading (SHER). The SHER project has been approved by the Swiss Association of Research Ethics Committees (CCER-2021-00528), and the parents signed the informed consent form before starting the data acquisition. All methods were carried out in accordance with Helsinki declaration.

Thirteen seven-month-old babies who were born premature were recruited by the Hopitaux Universitaires of Geneva. They took part in a two-month intervention program based on weekly shared reading activities with the objective of enhancing infant development and parent-infant emotional co-regulation, in an easily implemented, time- and cost-effective early intervention program. Families had to perform a minimum of 10 minutes per day four times per week of shared reading activity, using an age-tailored set of books and following specific strategies for effective shared reading. In the control group nine preterms spent an equal amount of time on shared play focused on spatial and building activities. Finally, an additional passive control group of fifteen babies born at term has been recruited at 9-months of age. More details on the participants of the three groups are reported in Table 1.

**Table 1:** Participant’s characteristics. GA stands for Gestational Age.

| Group | N. of infants | Female | Age at testing<br>[mean ± std] | GA at birth<br>[mean ± std] | Mother’s age<br>[mean ± std] |
| --- | --- | --- | --- | --- | --- |
| **Term** | 11 | 7 | 9 months 14 days<br>± 20 days | / | / |
| **Intervention** | 12 | 5 | 9 months 29 days<br>± 27 days | 34 weeks 1 days<br>± 6 days | 34.82 years<br>± 6.55 years |
| **Control** | 8 | 4 | 10 months 11 days<br>± 11 days | 33 weeks 6 days<br>± 4 days | 37.75 years<br>± 4.13 years |

Physiological and behavioural data of each parent-baby dyad were collected to assess the short- and long-term effects of the early intervention. The experimental protocol used to acquire physiological data was the same for the three groups (intervention, control, and term). The experimental sessions were conducted in an isolated space where standard play and reading kits were provided, and each dyad could interact in the most intimate and natural way possible. The experimenters observed the progress of the protocol from an adjacent space without being seen by the participants and interfering as little as possible with their activities. During the experimental protocol parents and infants performed three minutes of shared play and three minutes of shared reading activity. Electrocardiographic (ECG) data of each dyad were acquired simultaneously during the protocol by using a BIOPAC MP160 system, with a samplig rate of 1000 Hz. The ECG signals were pre-processed using Kubios 2.2 to detect R-peaks and correct for artefacts and ectopic beats. Six subjects were discarded due to low SNR, for a total of 12 preterm dyads in the intervention group, 8 preterm dyads in the control group and 11 term dyads as passive control. Then, we extracted the HRV signals after a piecewise cubic spline interpolation at 4 Hz.

The influence of the respiratory components was removed by combining the techniques reported in (41; 42; 43). Firstly, a respiratory signal (ECG-DR) was derived from the corresponding ECG using a slope-range method (43). The slope range is defined as the difference between maximum up-slope and minimum down-slope within the QRS interval, without considering their relative time of occurrence. The derived signals were post-processed by performing an outlier correction procedure according to (44), and sampled at 4 Hz using a cubic spline interpolation method. Afterwards, the orthogonal subspace projection method (41) was used to decompose the HRV in a respiratory component and a residual component containing sympathetic modulations and nonlinear respiratory interactions. The alorithm creates a subspace representing the respiratory activity, defined using the ECG-DR. The original HRV is projected in the subspace to find the component linearly related to respiration, while the residual component is defined as the orthogonal ones, and was used for the analysis, after z-score normalization. The MCI and *op*MCI indices were used to discriminate between different levels of coupling between the different groups and during the different tasks.

#### Statistical analysis

Concerning the synthetic dataset, we tested statistical differences in short scales (MCI*_short_* and *op*MCI*_short_, β* ∈ [1, 5]), long scales (MCI*_long_* and *op*MCI*_long_, β* ∈ [6, 10]) and total range of scales (MCI*_tot_* and *op*MCI*_tot_*), between the different level of *a_n_* in each configuration (unidirectional and bidirectional coupling). The normality of data distributions of the resulting features was assessed using the Shapiro-Wilk test, afterwards a one-way ANOVA was performed comparing the different groups, one for each value of *a_n_*, in each of the two configurations separately. The Bonferroni method was used to perform the p-value correction after the multiple comparisons tests.

When we applied statistical analysis to physiological data, we tested the differences between MCI*_short_*, MCI*_long_* and MCI*_tot_* (and between *op*MCI*_short_, op*MCI*_long_* and *op*MCI*_tot_*) among the three groups (term, intervention, control) and experimental sessions (play and reading tasks). The normality of the distribution of the resulting features was assessed using the Shapiro-Wilk test. Afterwards a two-way mixed ANOVA design was performed, with the group as the between subject factor, and the performed tasks as the within subject factor. The t-test was used as post-hoc test, paired for the task at each group level and unpaired for the group during each task level. Bonferroni’s correction was used for multiple comparisons.

## Results

### Synthetic data

Figure 1 shows the values of MCI obtained using the standard procedure in the unidirectional coupling and bidirectional coupling. Visibly higher values were reported in the case corresponding to higher coupling parameters, and statistically significant differences between the coupling levels were found for the indexes MCI*_short_*,MCI*_long_*, and MCI*_tot_* (*p <* 0.01 for all the significant comparisons). Concerning the unidirectional coupling case the post-hoc test showed that only the comparison between *a_n_* = 0 and *a_n_* = 0.02 at short term-scales was non-significant (*p >* 0.05). As regards the bidirectional coupling case the post-hoc statistical tests gave non-significant results for *a_n_* = 0 and *a_n_* = 0.02, at both short term- and long-term scales.

**Figure 1:**
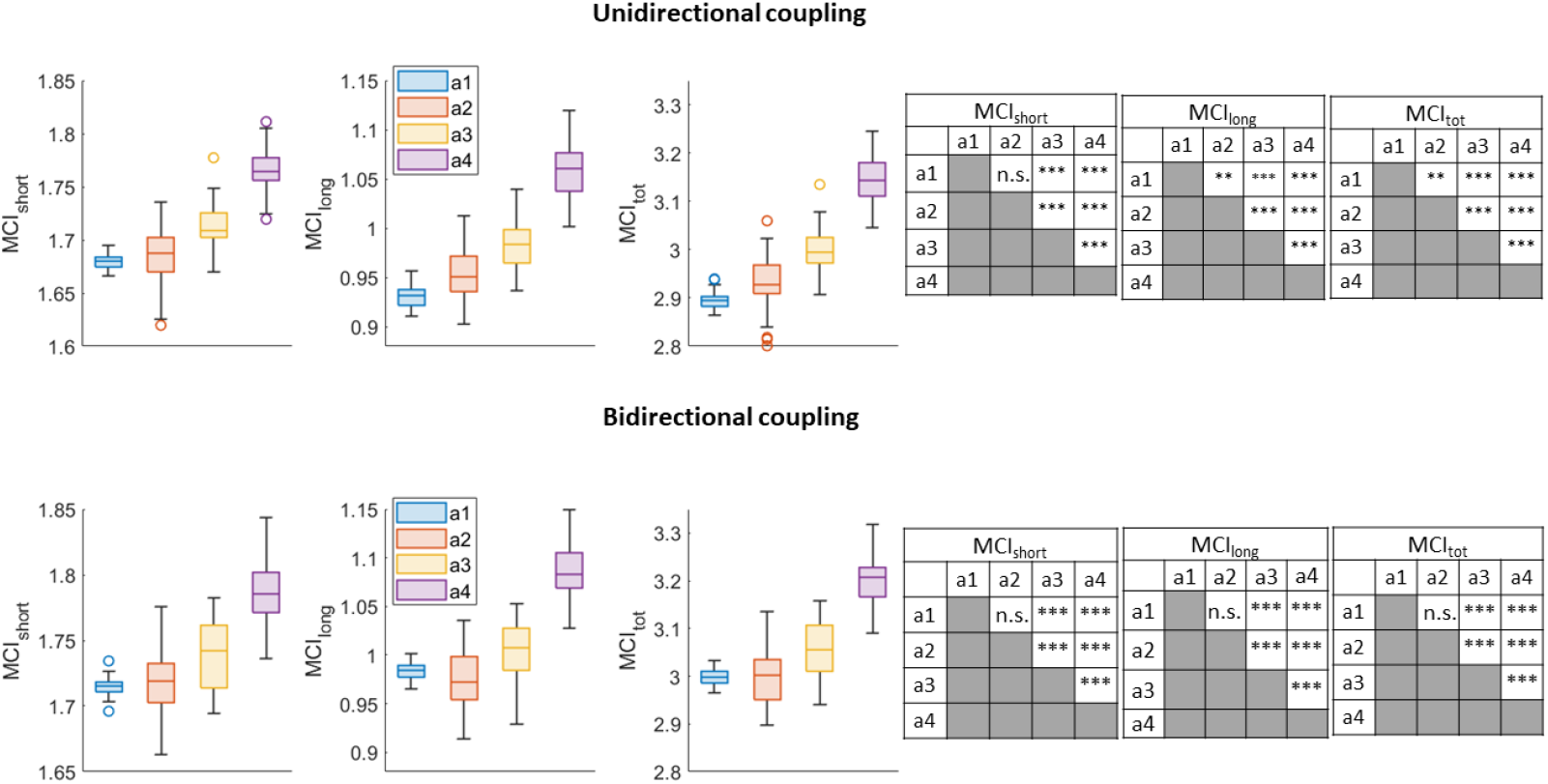
Results of the statistical analysis applied to MCI indexes on synthetic data generated by using the following values of coupling parameter *a_n_*: [a1 =0; a2=0.02; a3=0.04; a4=0.08]. The boxplots of the MCI values obtained from the 50 realizations for each *a_n_* value in the unidirectional coupling case (*b_n_* = 0) are shown on the upper left. *MCI_short_* corresponds to the scales *β* ∈ [1, 5], *MCI_long_* corresponds to the scales *β* ∈ [6, 10], while *MCI_tot_* corresponds to the area under the entire scale range. On the upper right, the tables summarize the significant differences in the comparisons between the *a_n_* values. Significance thresholds are represented by: * = *p <* 0.05; * * = *p <* 0.01; * * * = *p <* 10^−4^; n.s.= not significant. The corresponding results obtained for the bidirectional case (*b_n_* = 0.03) are shown on the lower part of the figure.

Figure 2 shows the results related to *op*MCI approach, after optimizing the MCI method by using the information of the cross-correlation. The trends obtained correspond with the previous results, showing higher *MCI* values in the cases of higher coupling parameters. The post-hoc tests showed that all comparisons between the different *a_n_* values were statistically significant (p-values in the range [0.02 − 10^−65^]) in both the unidirectional and bidirectional cases.

**Figure 2:**
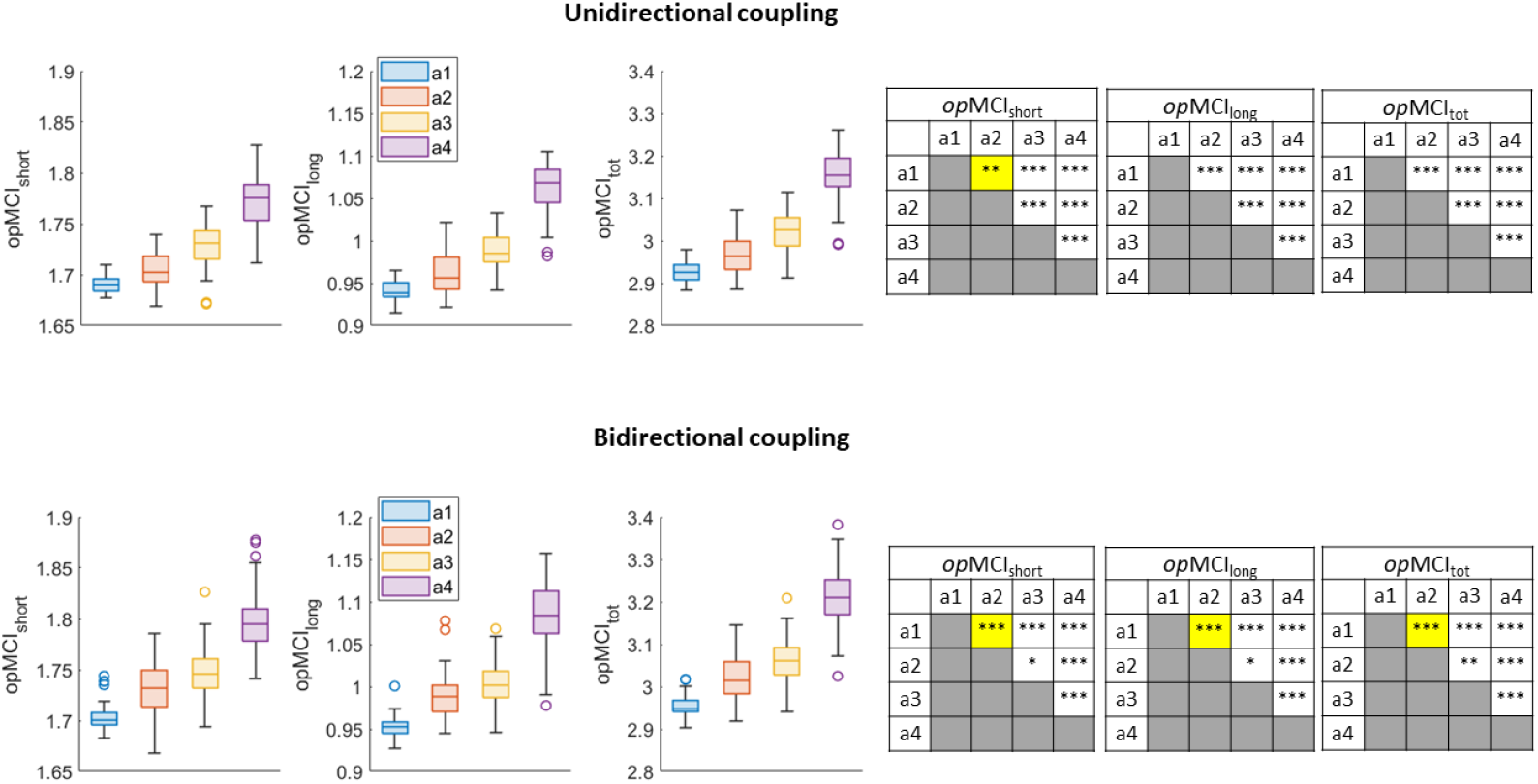
Results of the statistical analysis applied to *op*MCI indexes on synthetic data generated by using the following values of coupling parameter *a_n_*: [*a*1 = 0; *a*2 = 0.02; *a*3 = 0.04; *a*4 = 0.08]. The boxplots of the *op*MCI values obtained from the 50 realizations for each *a_n_* value in the unidirectional coupling case (*b_n_* = 0) are shown on the upper left. *op*MCI*_short_* corresponds to the scales *β* ∈ [1, 5], *op*MCI*_long_* corresponds to the scales *β* ∈ [6, 10], while *op*MCI*_tot_* corresponds to the area under the entire scale range. On the upper right, the tables summarize the significant differences in the comparisons between the *a_n_* values. Significance thresholds are represented by: * = *p <* 0.05; * * = *p <* 0.01; * * * = *p <* 10^−4^. The cells highlighted in yellow represent the additional significant differences with respect to the standard MCI index. The corresponding results obtained for the bidirectional case (*b_n_* = 0.03) are shown on the lower part of the figure.

The differences in values obtained between MCI and *op*MCI at each coupling level and scale range (short, long and total) have been assessed using a t-test, correcting afterwards with Bonferroni’s correction for multiple comparisons. Only comparisons between MCI and *op*MCI in the uncoupled realizations (a*_n_* = 0) of both bidirectional and unidirectional coupling cases displayed significant differences (*p <* 10^−4^) at all scales. At very low coupling (a*_n_* = 0.02), instead, the difference was significant (*p <* 0.01) only at short scales (*β* ∈ [1, 5]) and over the total scale range (*β* ∈ [1, 10]) for both unidirectional and bidirectional coupling. With medium coupling (a*_n_* = 0.04) a significant difference (*p <* 0.01) was found only in the unidirectional case at short scale range (*β* ∈ [1, 5]). All other comparisons were nonsignificant.

### Physiological data

Figure 3 shows the statistical results related to the MCI indexes extracted from parent-infant physiological data. The inter-group analysis revealed that during both playing and reading the preterm infant-parent dyads in the intervention group displayed higher average values with respect to the other two groups, with the difference being statistically significant only in the shared reading. Specifically, the MCI values obtained at short scales for the reading phase showed a statistically relevant (*p <* 0.05) difference when the intervention group was compared to the control group. At long scales, while the trend was maintained, the significance of the difference between the control and the intervention group was lost after Bonferroni’s correction of p-values. Considering the whole scale range for the computation of MCI (MCI*_tot_*) the intervention group reported significant higher values with respect to the control group (*p <* 0.01).

**Figure 3:**
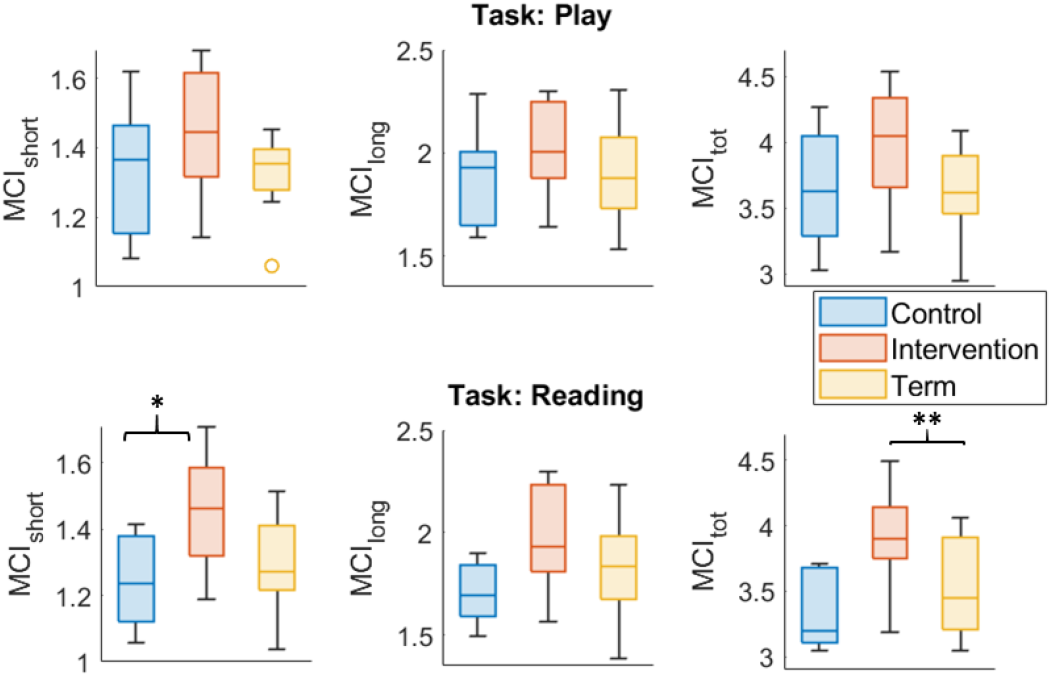
Boxplots related to the MCI values obtained from the parent-infant dyads’ HRV data using the standard approach. The MCI values of the three experimental groups (control, intervention, term) are compared during the play task (on the top) and during the reading task (on the bottom). MCI*_short_* corresponds to the scales *β* ∈ [1, 5], MCI*_long_* corresponds to the scales *β* ∈ [6, 10], while MCI*_tot_* corresponds to the area under the entire scale range. Significance thresholds are represented by: * = *p <* 0.05; ** = *p <* 0.01.

Figure 4 displays the results obtained using the proposed *op*MCI approach. Significant inter-group differences were found looking at the overall scale range during the reading phase: the intervention group showed significantly higher values of synchrony (*p <* 0.05) with respect to both control and term groups. Only the statistical significance of the differences between the intervention and control groups remained after Bonferroni’s correction on short-term scales.

**Figure 4:**
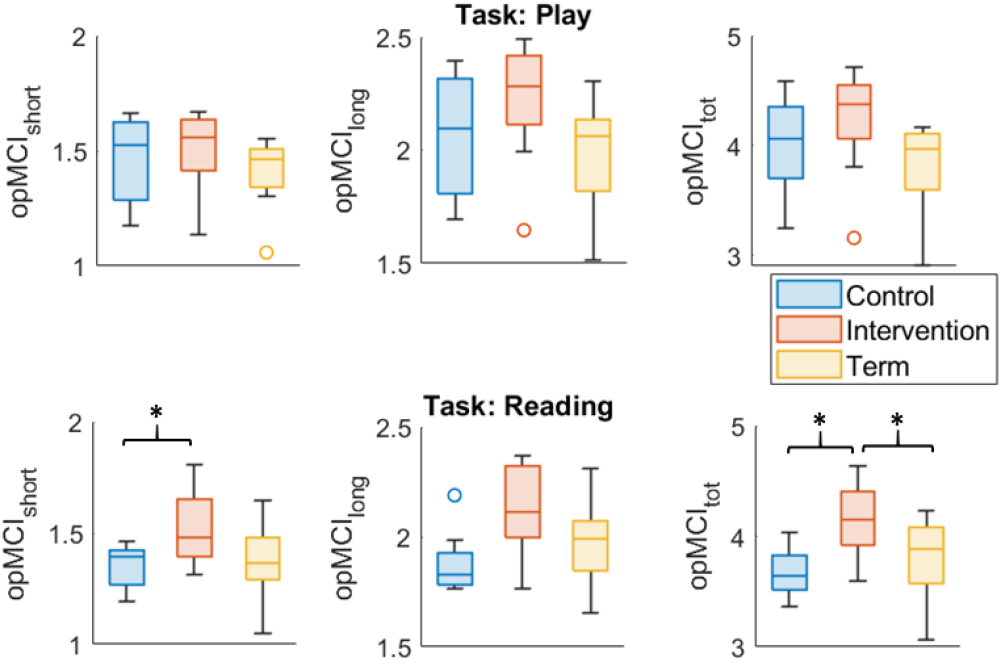
Boxplots of the *op*MCI values obtained from the parent-infant dyads’ HRV data. The three groups (control, intervention, term) are compared during the play task (on the top) and during the reading task (on the bottom). *op*MCI*_short_* corresponds to the scales *β* ∈ [1, 5], *op*MCI*_long_* corresponds to the scales *β* ∈ [6, 10], while *op*MCI*_tot_* corresponds to the area under the entire scale range. Significance thresholds are represented by: * = *p <* 0.05; ** = *p <* 0.01.

## Discussion

We presented *op*MCI, a new approach to assess physiological synchronization of different individuals, focusing on the dynamical nature of social interactions and on the intrinsic complexity of multiscale psychophysiological phenomena. The *op*MCI method is an optimization of the MCI index proposed by Nardelli *at al*. in (34). The capabilities of *op*MCI to assess the coupling between dynamical systems were tested on synthetic time series taken from a system of two Hénon maps with four different coupling levels (a_1_=0, a_2_=0.02, a_3_=0.04, a_4_=0.08), and on cardiovascular data of 31 parent-infant dyads acquired during a clinical experimental trial. For both synthetic and real data, we applied statistical tests in order to compare the performance of both *op*MCI and MCI indexes to discern coupling levels.

Concerning the application of MCI approach to coupled Hénon maps, significantly higher values were displayed when the coupling coefficient increased, allowing to statistically discern between the coupling levels in the case of both unidirectional and bidirectional couplings (see Figure 1). However, after the post-hoc analysis, the comparison between the smaller coupling coefficient *a_n_* = 0.02 and the uncoupled case *a_n_* = 0 was not statistically significant at all scale ranges for the bidirectional coupling and at the short scales for the unidirectional one. In the case of physiological data, a statistically significant difference was found between the preterm intervention and control group during the reading task (see Figure 3), for short scales (MCI*_short_*) and considering all the range of scales (MCI*_tot_*). The newly introduced optimization procedure allowed to further improve the performance of the MCI, taking into account the cross-correlation of the physiological series involved. As shown in Figure 2, the *op*MCI was able to discriminate the coupling value a*_n_*=0 from a*_n_*=0.02, correctly identifying all the different coupling levels for both the unidirectional and bidirectional case. Concerning the physiological data, an additional significant difference was found between the intervention and the term groups during the reading task at the full range of scales *op*MCI*_tot_* (see Figure 4).

This last promising result indicates that not only the intervention program increased synchrony in preterm dyads during the reading task, but that this synchrony displays higher values than dyads at term. The here proposed approach, based on multiscale dynamical interaction between biological systems, can be generalized to several different social contexts and clinical applications, investigating the nuances of synchrony between individuals by using ANS, interbrain and hormonal data (33).

In this study we applied our approach to HRV signals, due to the specific clinical and psychological relevancy of ANS in the dyadic interactions, and considering that it’s a well-validated tool for the recognition of emotions and physiological arousal states (13; 22; 34; 27; 30; 29). From a physiological point of view the combined activity of the two main branches, the Sympathetic Nervous System (SNS) and the Parasympathetic Nervous System (PNS), can offer valuable insights in inter-individual physiological synchronization. According to the literature, PNS activity can be involved in dyad’s ability to positively engage and flexibly respond to environmental changes (1). The SNS activity can be used as an indicator of the reciprocal role of dyads’ in modulating one another’s arousal, reducing or increasing, and in mutual stress responses (1; 45). From a technical point of view, the ANS activity characterization through HRV analysis allows ecological monitoring using non-invasive and wearable sensors (46). For this reason, the proposed approach is well suited for applications that rely on observing naturalistic interactions between individuals, which would be affected by the use of cumbersome instrumentation (47).

A limitation of the reported clinical application can be represented by the sample size. However, research involving infant data, encompassing the age range from birth to 12 months, frequently encounters the constraint of a relatively modest sample size. Challenges inherent in collecting data from infants include ethical considerations, consent procedures, parental involvement, logistical hurdles, and specific cultural needs. Notably, our focus on preterm-born infants adds an additional layer of complexity to recruitment post-hospitalization, exacerbated by their propensity for recurrent health issues, such as seasonal infections, during development.

The causality of the interaction will be investigated in future studies to assess the uni- or bidirectionality of coupling and provide key insights regarding the roles of the dyad participants in the social task. In cross-sectional studies on parent-infant synchrony the direction of coupling could be a relevant factor for the task being performed (1). Future longitudinal studies are required to disentangle the physiological changes in infants becoming more socially responsive and active during the interactions, and how this could be affected by premature birth (17; 19; 16). In addition, the *op*MCI application could be extended from dyad to group social interactions, further enlarging the application of this approach to non-clinical fields, e.g., education, sport, team-building (2; 13).

